# The effects of fixational tremor on the retinal image

**DOI:** 10.1101/403964

**Authors:** Norick R. Bowers, Alexandra E. Boehm, Austin Roorda

## Abstract

The study of fixational eye motion (FEM) has implications for the neural and computational underpinnings of vision. One component of FEM is tremor, a high-frequency oscillatory jitter reported to be anywhere from ∼5 to 60 seconds of arc in amplitude. In order to isolate the effects of tremor on the retinal image directly and in the absence of optical blur, high-frequency, high-resolution eye traces were collected in 6 subjects from videos recorded with an Adaptive Optics Scanning Laser Ophthalmoscope. Videos were acquired while subjects engaged in an active fixation task where they fixated on a tumbling E stimulus and reported changes in its orientation. Spectral analysis was conducted on isolated segments of optical drift. The resultant amplitude spectra showed a slight deviation from the traditional 1/f nature of optical drift in the frequency range of 50-100 Hz, which is indicative of tremor; however, the amplitude of this deviation rarely exceeded one second of arc, smaller than any magnitude previously reported.

## Introduction

### Fixational Eye Motion

Even during intersaccadic periods of fixation the eye is never still; small eye movements constantly shift the retinal image over the photoreceptor mosaic (for review, see Martinez-Conde, Macknik, & Hubel, 2004). These movements can shift a stimulus over dozens of photoreceptors every second. Fixational eye motion has typically been categorized to fall into three main components; (i) Microsaccades, small ballistic movements similar to larger saccades, (ii) Ocular Drift, a slow Brownian-like movement shifting the gaze only a few arcmin, and (iii) Tremor, a high-frequency oscillatory jitter roughly the size of a foveal cone (Ditchburn & Ginsborg, 1953; Eizenman, Hallett, & Frecker, 1985; Ko, Snodderly, & Poletti, 2016; Ratliff & Riggs, 1950; Rucci & Poletti, 2015). Extensive research has been conducted on the functional and perceptual consequences of microsaccades and drift (Bowers & Poletti, 2017; Burak, Rokni, Meister, & Sompolinsky, 2010; Kagan, Gur, & Snodderly, 2008; Ko, Poletti, & Rucci, 2010; Kuang, Poletti, Victor, & Rucci, 2012; Ratnam, Domdei, Harmening, & Roorda, 2017; Rucci, Iovin, Poletti, & Santini, 2007), but the perceptual consequences of tremor and its possible functional role are still not fully understood.

### Tremor

Reports of tremor vary widely on the statistical nature and magnitude of this motion. Tremor is generally defined as an increase in eye motion amplitude at high frequencies. The bandwidth of tremor is often reported as falling between 50 Hz and 100 Hz, whereas the amplitude has been found to be as small as 4.8 arcseconds (Ko, et al., 2016) and as large as 1 arcminute (Ratliff & Riggs, 1950) and some reports question the existence of tremor at all (Stevenson, Roorda, & Kumar, 2010). Few studies have set out to examine the implications of tremor for vision, largely due to the technical difficulties of accurately measuring tremor with conventional eye trackers. There is some evidence that tremor could contribute to perception by synchronizing retinal ganglion cells (Greschner, Bongard, Rujan, & Ammermüller, 2002) or through stochastic resonance of visual noise (Hennig, Kerscher, Funke, & Wörgötter, 2002).

### Eye Tracking

Most reports of tremor stem from the use of high-resolution eye tracking techniques, such as dual-Purkinje image (DPI) eye tracking (Crane & Steele, 1985; Ko et al., 2016), scleral search coils (Houben, Goumans, & Van Der Steen, 2006), reflections from small mirrors placed on contact lenses (Ditchburn & Ginsborg, 1953; Ratliff & Riggs, 1950; Riggs & Schick, 1968; Steinman, Haddad, Skavenski, & Wyman, 1973; Yarbus, 1968) and reflections from the cornea directly (Eizenman et al., 1985). Each of these trackers has the potential spatial and temporal resolution to measure tremor, but each relies on tracking some part of the anterior segment or lens of the eye and inferring the motion on the retina. The current study looks to examine the effects of tremor on the retinal image directly using an Adaptive Optics Scanning Laser Ophthalmoscope (AOSLO), a relatively novel method of tracking the eye that relies on imaging the retinal surface directly (Stevenson & Roorda, 2005; Vogel, Arathorn, Roorda, & Parker, 2006).

## Methods

### AO System

Movies of the retina are obtained through the use of an Adaptive Optics Scanning Laser Ophthalmoscope (AOSLO) (Roorda et al., 2002). In the AOSLO system, a focused point of light is scanned across the retina in a raster pattern to obtain high-resolution movies of the photoreceptor mosaic during fixation. In the most recent version of the system, a supercontinuum light source provides a point source of light that is relayed through the optical path to a fast horizontal resonant scanner (16 kHz) and a slow vertical scanner (30 Hz), which together sweep the point across the retina in a raster pattern. and the deformable mirror (7.2 mm diameter 97 actuator membrane; ALPAO, Montbonnot-Saint-Martin, FRANCE), which compensates for the aberrations of the eye. The light reflecting from the retina is relayed and descanned back through the optical path to a custom Shack-Hartmann wavefront sensor, which measures the aberrations, and through a confocal pinhole (conjugate to the retinal plane of focus) and then to a photomultiplier tube, which is used to record the scattered light, pixel-by-pixel, to reconstruct an image of the retinal surface. The AOSLO system used is equipped with 4 wavelength channels: 840, 680 and 543 nm channels are used for imaging and stimulus delivery and a 940 nm wavelength channel is used for wavefront sensing. In this particular experiment, 840 nm was used for imaging and 543 nm was used to provide a stimulus for fixation (see Experimental Design). The vergence of all wavelengths were adjusted in the light delivery arm to compensate for longitudinal chromatic aberration so that all wavelengths were in simultaneous focus on the retina (Harmening, Tiruveedhula, Roorda, & Sincich, 2012; Grieve, Tiruveedhula, Zhang, & Roorda, 2006). Custom software was used to operate the entire AO control system. Measurement and correction were performed over the entire pupil diameter up to a maximum of 7.2 mm. This system generally obtains near diffraction-limited images of the retina with high enough spatial resolution to resolve foveal cones.

### Strip-Based Eye Tracking

The images obtained by the AOSLO system were compiled together in a continuous sequence to create movies of the retina. The movies were acquired at 30 Hz (the frequency of the slow vertical scanner) and were composed of 512×512 pixels. The size of the raster on the retina was 0.9 degrees so that each arcmin is subtended by ∼10 pixels. Eye movement traces were acquired from the movies using an offline algorithm that utilized a strip-based cross correlation technique to obtain eye traces at higher temporal resolution than the frame rate of the system (Stevenson & Roorda, 2005; Vogel, Arathorn, Roorda, & Parker, 2006). Since each frame was acquired over time, additional temporal information on eye movements, which manifest as unique distortions within each frame, is available beyond the 30 Hz frame rate. The top of each frame occurs earlier in time than the bottom, and by dividing each frame into strips and analyzing the movement in a strip-wise manner, eye motion traces can be acquired with a much higher temporal sampling rate than the frame rate of the movie (30 Hz). The eye motion sampling rate is the frame rate multiplied by the number of strips per frame, so the temporal resolution of the eye motion can be adjusted by increasing or decreasing the number of strips used in the cross-correlation. For the current study 64 strips were used per frame, giving an eye motion sampling rate of 1920 Hz. The eye motion correction has been done in real time (Arathorn et al., 2007) and offline (Stevenson et al., 2010). For all analyses in this paper, eye motion computations were done offline after the videos were acquired.

For offline analysis, an oversized composite reference frame is generated for each movie by averaging together and roughly aligning selected frames of the movie. The size of the composite reference is dependent on the extent of eye motion during recording. Each frame of the movie is then divided into 64 strips that are 8 pixels in height and run the entire 512 pixels of the frame width. Each strip is cross-correlated against the reference frame in order to obtain vertical and horizontal offsets of the eye position compared to the reference at that instance. Each strip represents 1 sample of the eye trace and the strips from each frame are strung together into a continuous sequence to obtain eye traces at a rate of 1920 Hz from the 30 Hz AOSLO movies. This technique is able to record eye motion with amplitudes smaller than one second of arc (Stevenson et al., 2010).

### Eye Movement Parsing

Once the raw eye traces were acquired, they were parsed using an automatic algorithm. First, erroneous or noisy eye motion traces recorded during blinks or during periods where the image quality was very poor were identified by labeling frames in the movie in which the average luminance of the total frame fell below a threshold. Second, saccades were identified using a speed threshold, where saccade onset was considered the point in which the eye moved above 1.5 deg/sec and saccade offset was considered the point in which the eye fell below 1.5 deg/sec. Saccades falling above an amplitude threshold (3 arcmin) were not considered for analysis. Drift was identified as all intersaccadic periods of eye motion. Saccade detection was verified by human observers manually to identify any saccades the automatic algorithm may have missed. In the data sets collected here on young healthy eyes with normal fixation (see Experimental Design), the automatic algorithm captured most saccades and only a small number had to be manually identified. Whenever a frame contained poor data or a saccade, the entire frame was flagged and not considered in the analysis. The first sample of each drift frame was repositioned to align with the last sample of the previous drift segment to eliminate discontinuous jumps from saccades and blinks. This technique was used because intersaccadic periods of drift are generally small (∼300 ms), which poses a constraint on the resolution of the Fourier analysis. Drifts stitched together in this method were cropped together using only full frames worth of samples (∼33 ms periods), that is, any portion of a drift that began or ended in the middle of a frame was not included. This was done in order to better eliminate periodic artifacts at the frame rate of the system arising from torsion or reference frame distortions (see Reference Frame & Torsion Correction). This method assumes stationarity of intersaccadic drift segments (disregarding any polar bias in drift direction).

### Reference Frame & Torsion Correction

Although the eye motion traces after parsing produce continuous segments of isolated drift at high sampling rates, the traces still contain motion artifacts caused by distortions in the reference frame as well as torsional eye movements. Torsional eye movements produce a sawtooth waveform that repeats at the frame rate of the system, whereas reference frame distortions present as a short random walk overlaid onto each frame’s motion. Fortunately, both of these artifacts are periodic and introduce peaks in the amplitude spectra that are isolated to the frame rate of the system and higher harmonics only; they do not affect the underlying amplitude spectrum anywhere else. In the eye motion traces from the offline processed videos, reference frame artifacts are largely removed by using multiple frames to generate a composite reference frame (Stevenson et al., 2010). By combining a series of frames, the distortions of the individual frames are largely averaged out (Bedggood & Metha, 2017). Torsion, however, may change over the course of a video and the periodic sawtooth must be removed from each frame individually. A full description of the algorithms to measure and remove reference frame distortion and torsional artifacts is part of a paper in progress. For the purposes of this paper, it is sufficient to state that the sawtooth artifact was measured and removed from the eye motion trace prior to further analysis.

### Eye Motion Analysis

The amplitude spectra of these drift segments were then analyzed using Fourier analysis, specifically using a method proposed by Welch (1967). This method involves dividing the signal into overlapping segments and calculating the amplitude spectrum for each segment which are then averaged together. Each sample was weighted in such a way that its multiple inclusion in the analysis is nullified and its overall contribution to the amplitude spectra was the same as any other sample. In order to maximize resolution, the window size selected was equal to the sampling rate of the eye trace (1920 Hz), allowing for analysis of frequencies up to 960 Hz. Outputs were converted to amplitude in arcminutes.

### Validations

In order to validate the ability of the AOSLO system to track eye motion, two simulations were conducted. The first simulation aimed to test the ability of the entire AOSLO system to record motion. A model eye was attached to a galvanometer and oscillated in a diagonal sinusoidal pattern at a variety of frequencies (4 Hz, 16 Hz, 64 Hz, and 256 Hz) while being recorded with the AOSLO system. To enable analysis of the noise floor of the system, a video was also recorded of the non-moving model eye. The results of this simulation can be seen in Figure 2. Thirty second movies were collected at each frequency with an amplitude of 0.45 arcmin at an angle of ∼26.6 degrees to give horizontal and vertical amplitudes of ∼0.4 and ∼0.2 arcmin respectively. The motion from these movies was processed using the same methods used for movies of human eye motion.

The second simulation was aimed at validating the ability of the strip-based cross correlation technique for recovering motion from actual AOSLO movies. First a real AOSLO movie was manipulated digitally to add a distortion consistent with independent sinusoidal motions in the vertical and horizontal directions with a fixed amplitude. Following the manipulation, the movie was analyzed using the strip-based cross correlation technique described above to obtain traces of eye motion at 1920 Hz. The amplitude spectra of the motion trace from the manipulated movie were compared against the amplitude spectra of the motion trace from the original movie. We felt that validation with an actual AOSLO video was important because in a model eye the luminance and contrast of the image is static and the retina moves in only the direction of the galvanometers. The AOSLO movie, by comparison, contains actual eye motion and has more variable luminance owing to change in the adaptive optics correction as well as actual changes in reflected intensity from the retina (Pallikaris, Williams, & Hofer, 2003). If our eye motion analysis can recover the frequency and amplitude of high frequency motion that has been added to an actual movie, then we can be confident that the strip-based eye tracking algorithm would be able to detect real eye motion at these frequencies.

### Experiment Design

Six healthy subjects with normal vision were recruited for the study. Informed consent was obtained from each subject and all experimental procedures were reviewed and approved by the UC Berkeley Institutional Review Board and adhered to the tenets of the Declaration of Helsinki. To prepare subjects for AOSLO imaging, one drop of tropicamide (1%) and phenylephrine (2.5%) solution were administered topically 15 minutes prior to imaging to temporarily dilate the pupil and paralyze accommodation. For measurements of the motion amplitude spectrum in human eyes, AOSLO videos were recorded while subjects fixated a letter E optotype (Figure 1). The E was projected directly onto the retina at the center of the raster scan using the green channel (543 nm) in the AOSLO system while imaging was done with 840 nm NIR light. The combination of NIR light and very weak green background light (caused by light leaking through the acousto-optic modulator (AOM) that was used to project the letter ‘E’), formed a dim, reddish background over the extent of the raster scan. The E was 5 minutes of arc in height, corresponding to a 20/20 letter. To keep the subject engaged in the fixation task over the course of each video, the E changed orientation randomly on average once a second and the subject was instructed to report its orientation every time it changed via the arrows on a keyboard. Each subject completed five 30-second trials, for a total of 2.5 minutes of fixational eye motion per subject (with the exception of one subject, who only had four 30-second trials).

**Figure 1.**
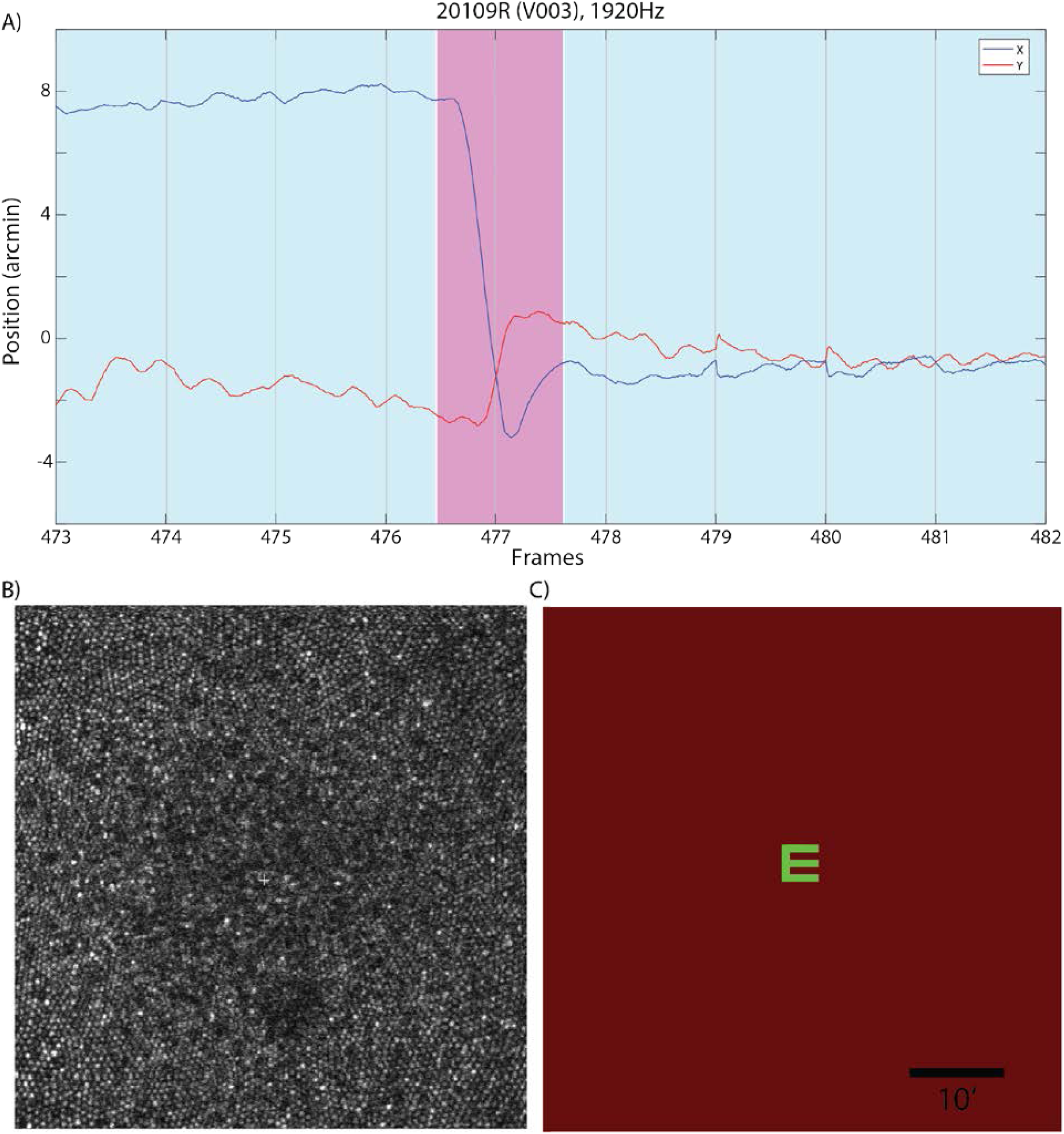
A) An example of an eye trace taken from an AOSLO movie. A microsaccade (magenta) is clearly distinguishable from the ocular drift (blue). Gray lines demarcate different frames. B) An example of an image/frame from the AOSLO system. The cone mosaic can be resolved even at the fovea. C) An example of the AOSLO raster with a green letter ‘E’ that subjects would see.

## Results

The first simulation aimed to test the entire AOSLO system’s ability to detect sinusoidal oscillations from a moving model eye. The resultant amplitude spectra plotted in Figure 2 showed a clear peak at the frequency of the sinusoidal oscillation for all input frequencies. Even though the motion of the sinusoidal oscillation was just a fraction of an arcmin, the AOSLO system was able to reliably recover the amplitude of the input motion. The slight reduction in measured amplitude at the higher frequencies resulted from the finite width of the sampling window (8 pixels) and small shifts in frequency of the galvanometer scanner over the course of the video.

**Figure 2.**
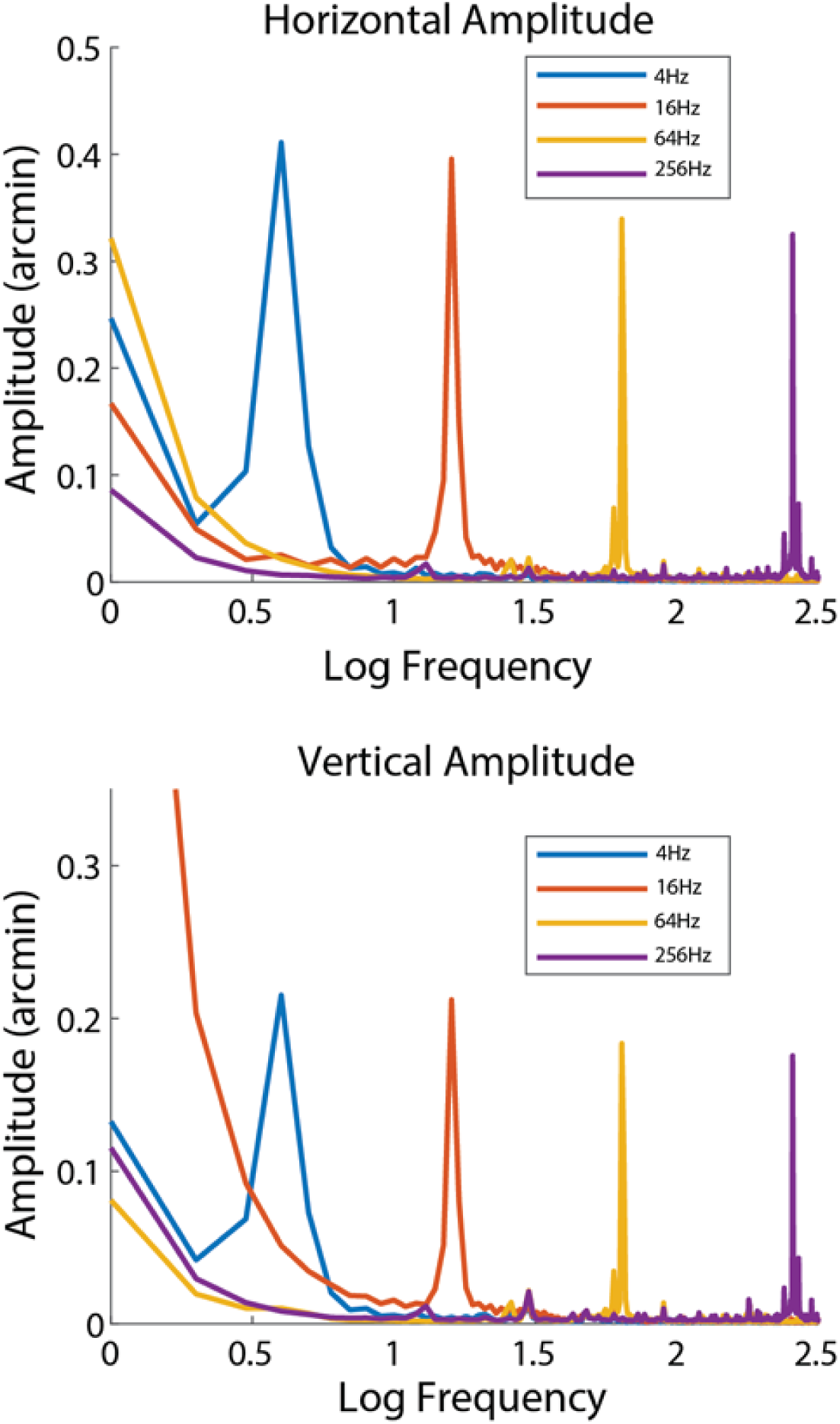
Recovered amplitudes for model eye movies. The frequencies and amplitudes input to the galvanometer scanner was readily recovered in the AOSLO movies of the model eye.

The second simulation aimed to test the capabilities of the strip-based cross-correlation technique to recover motion from AOSLO movies. A real AOSLO movie was manipulated to add distortions consistent with a sinusoidal motion of two frequencies – a horizontal frequency of 75 Hz and a vertical frequency of 50 Hz – both with an amplitude of 6 arcseconds. The distorted movie was then processed using the same strip-based cross correlation technique used on the human eye motion. The resultant amplitude spectrum of the manipulated movie compared to the original movie plotted in Figure 3 showed strong correlation except for a large peak in the manipulated movie’s horizontal spectrum at 75 Hz and the vertical amplitude spectrum at 50 Hz. Note that the amplitude spectra should theoretically match perfectly at all other frequencies, however the inclusion of the sinusoidal oscillation subtly changed the samples which were flagged as saccades, so some small discrepancy between the two is to be expected. Regardless of this discrepancy, the strip-based cross correlation technique was robustly able to recove the exact frequency and amplitude of the sinusoidal oscillation added.

**Figure 3.**
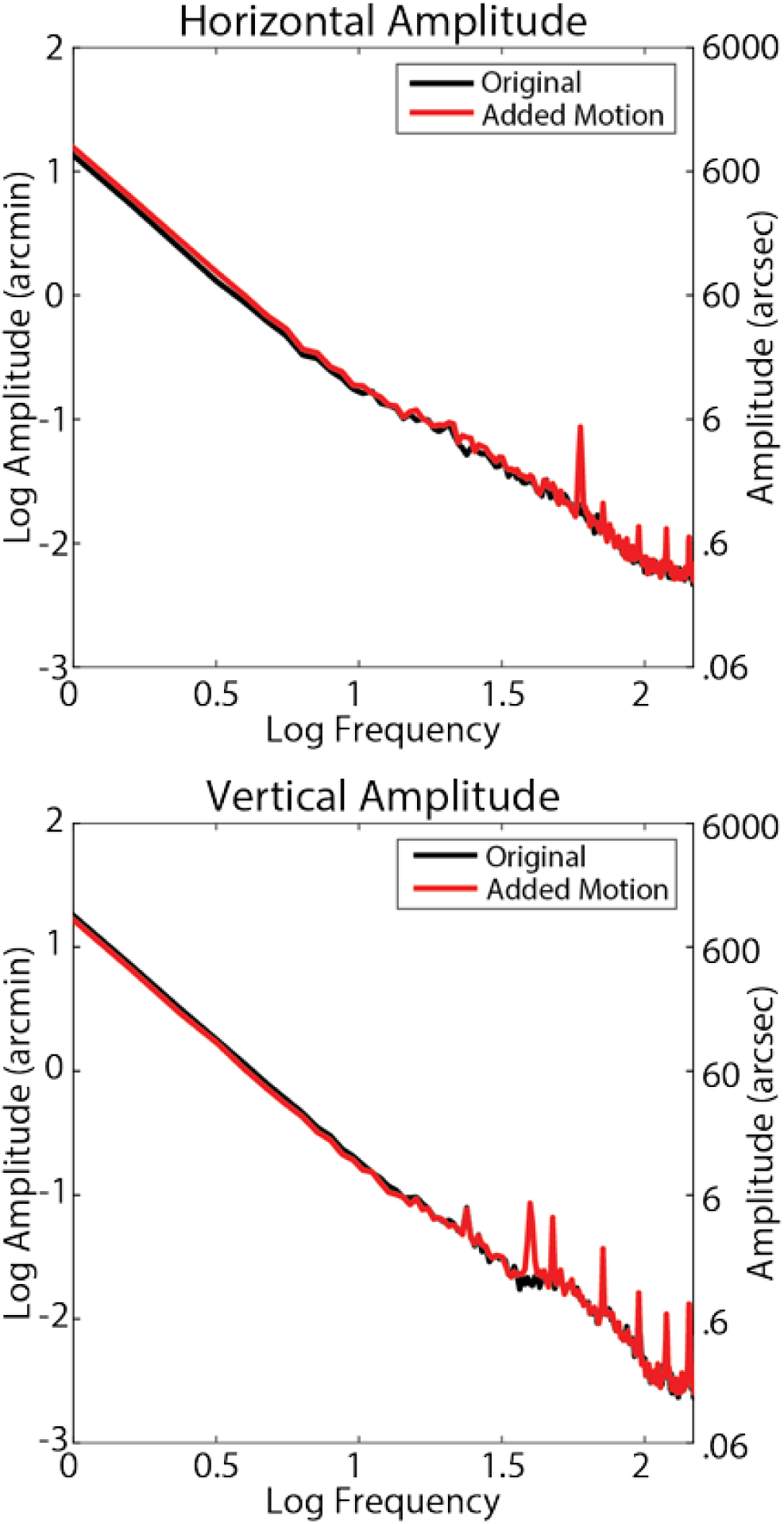
Amplitude spectra of original AOSLO movie (black) and modified AOSLO movie (red). The 6’ peak in the amplitude spectra at 75 Hz that was artificially added to the horizontal motion is clearly visible. In the vertical amplitude spectra, a 6’ peak was artificially added at 50 Hz. The peaks at higher frequencies, especially visible in the vertical dimension, are harmonics of the frame rate (90, 120, 150 Hz) from torsion that were not fully removed.

Using the technique outlined above, amplitude spectra of fixational drift for N=6 subjects were calculated. Heat maps for saccade landing positions and drift are shown on Figure 4. All subjects showed normal fixational eye motion. Subjects made microsaccades roughly once per second (1.10 ± 0.57 microsaccades/sec) with a normal amplitude (7.5’ ± 1.5’), speed (375 ± 49 arcmin/sec), and duration (36.9 ± 6.9 ms). Intersaccadic periods of drift also showed relatively normal statistics. Amplitude (3.8’ ± 0.9’), span, defined as the maximum distance from the mean location (3.2’ ± 0.7’), speed (79.6 ± 15.6 arcmin/second) and duration (620 ± 245 ms) were all within normal parameters and were consistent with previous findings ((Martinez-Conde, Macknik, & Hubel, 2004; Rucci & Poletti, 2015). Overall subjects did show slightly better fixational stability than average. This is likely due to all subjects being highly trained in psychophysics experiments using the AOSLO system.

**Figure 4.**
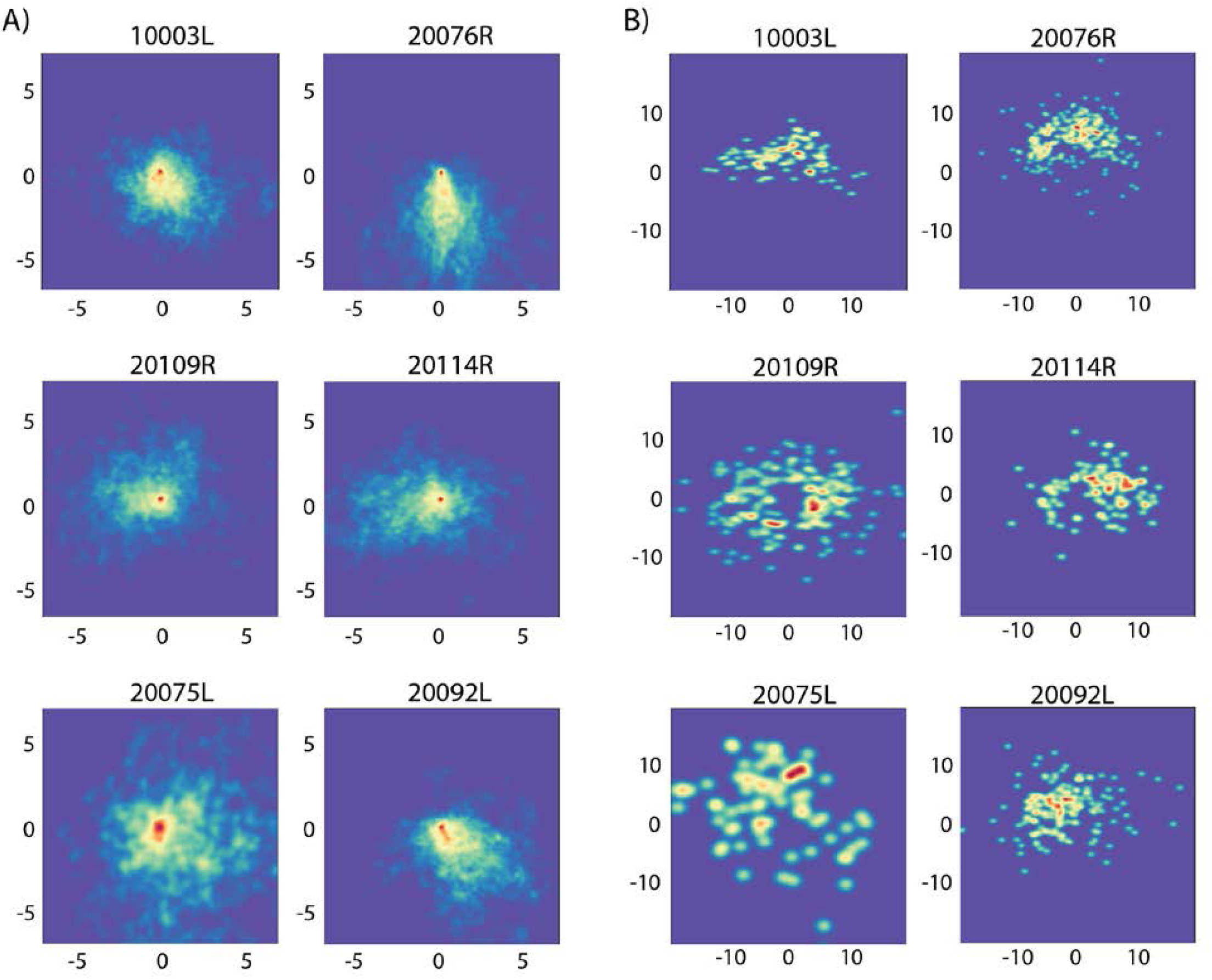
Heatmaps of position and amplitude for drift (A) and saccades (B). All data points are normalized at 0 for display purposes. All units are in minutes of arc. Each panel is normalized based on the amount of data available for that subject.

The average horizontal and vertical amplitude spectra for all 6 subjects are shown in Figure 5. Similar to that reported in Ko et al (2016), the spectra show a steeper than 1/f fall-off in amplitude, but becomes 1/f after 10 Hz. However, unlike Ko et al and others who used different tracking methods, the characteristic deviation from 1/f in the amplitude spectra indicative of tremor was not readily observed. Although there was a slight elevation within the band of 50 Hz – 100 Hz, the amplitude of this deviation rarely exceeded ∼1 arcsecond, much smaller than previous reports of tremor (∼30’) (Ko, et al., 2016; Ratliff & Riggs, 1950).

**Figure 5.**
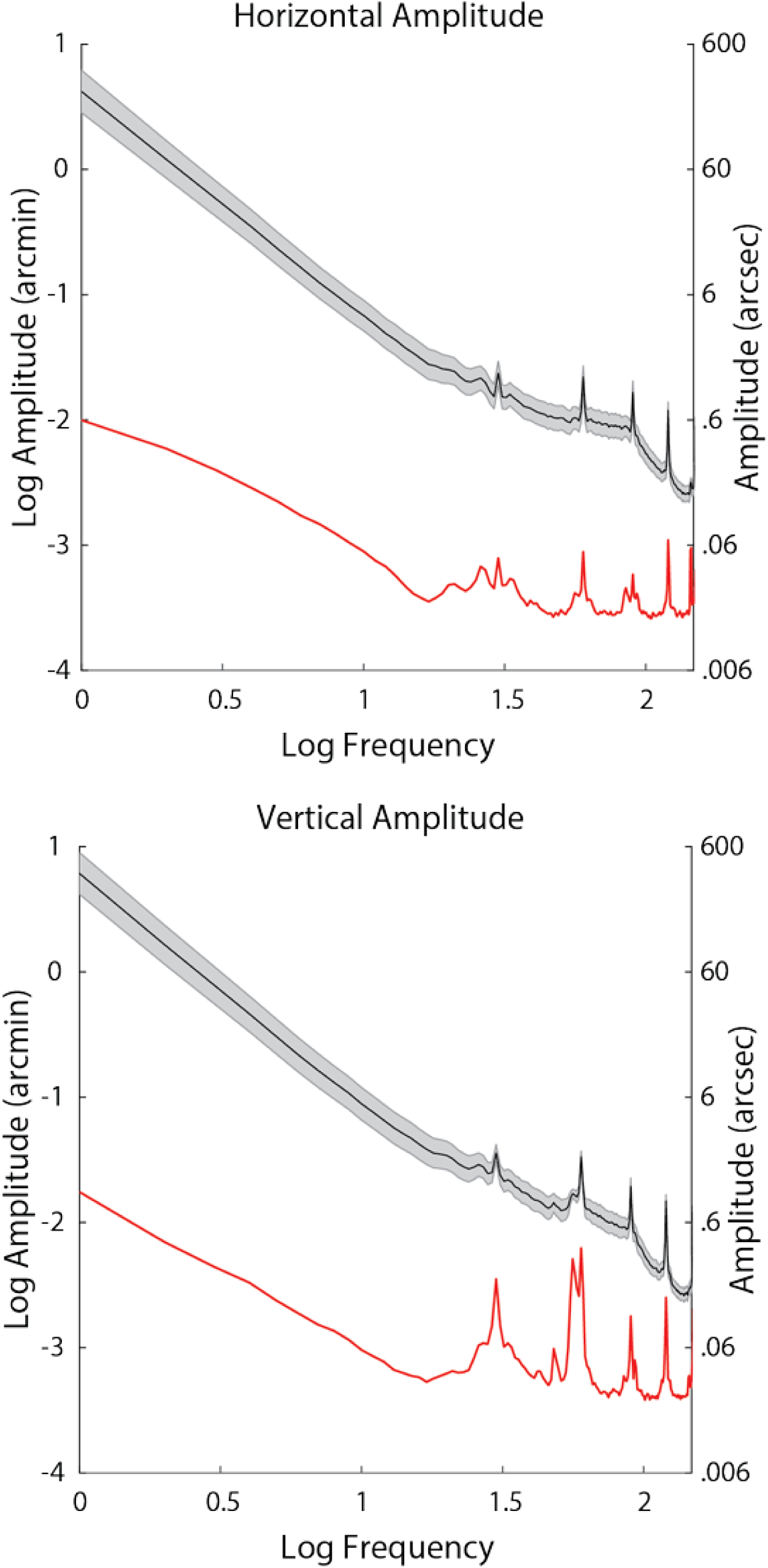
Mean amplitude spectra for 6 subjects (black) and noise floor measured from a non-moving model eye (red). The slight deviation between 50-100 Hz indicative of tremor is less than 1 arcsecond, much smaller than all previous reports of tremor. Shaded error bars represent ± 1 S.E.M. The spikes in the spectra are from periodic artifacts in the traces caused by tremor and residual reference frame distortions.

## Discussion

In this paper we validate and demonstrate the use of AOSLO as an eye tracker capable of recording microscopic eye movements with high fidelity up to very high frequencies. The temporal sampling rate of eye traces from the AOSLO system (1920 Hz) allows for analysis of frequencies up to 960 Hz, well beyond the 50 Hz-100 Hz bandwidth of tremor. The noise floor in the 50-100 Hz range measured from a non-moving model eye is ∼0.2 arcsec, which is well below the amplitude of any eye motion previously reported, including tremor. The eye tracking capabilities of the AOSLO system, combined with direct retinal imaging, is therefore uniquely capable of analyzing the effects of small eye movements on the retinal image. We validated the technique by recovering motion from a model eye that moved with specific frequencies and amplitudes and also by recovering motion that was digitally added to actual AOSLO videos. In both cases the AOSLO system was effective in recovering the motion.

The measurements of retinal image motion from the 6 normal subjects during active fixation showed some evidence of tremor in the frequency range of 50 to 100 Hz, but nothing with an amplitude greater than 1 arcsecond. Even when there was evidence of tremor, the amplitude power spectrum was monotonic (continuously declining) with increasing frequencies and was little more than a slight deviation from a 1/f curve.

Although there are some suggestions tremor could contribute to the visual percept through synchronization of retinal ganglion cells or through influencing the behavior of visual neurons in the brain (Greschner et al., 2002; Hennig et al., 2002), these studies generally assume the amplitude of tremor is around the scale of a foveal cone. Given that tremor on the retina is much lower than this, the possibility of this movement influencing the visual percept will need to be reexamined.

### Effects of Cycloplegia

In the current study, it was necessary to dilate and cycloplege subjects’ eyes in order to achieve the best image quality for image-based eye tracking. Cycloplegia relaxes the ciliary muscle but, being that it is a sphincter muscle, it actually leads to an increase in the tension on the lens. Decreasing the tension on the lens is known to increase lens wobble (He, Donnelly, Stevenson, & Glasser, 2010). How this intervention might affect the magnitude and or presence of tremor is not well known. To address this question, we performed similar eye-tracking measurements in a tracking scanning laser ophthalmoscope (for details on that system, see Sheehy et al (2012) for subjects that had not been cyclopleged. One 1-minute video was collected for 6 subjects (four of whom also participated in the previous experiment) and the amplitude spectra was assessed using the same method described above. The results are shown on Figure 6. Compared to AOSLO, the sampling resolution was ∼ 5 times lower (0.5 arcmin per pixel) and the image resolution was not as high, Nevertheless, the frequency bandwidth was the same and the noise floor was sufficient to detect tremor. Amplitude spectra measures without cycloplegia showed a similar lack of tremor.

**Figure 6.**
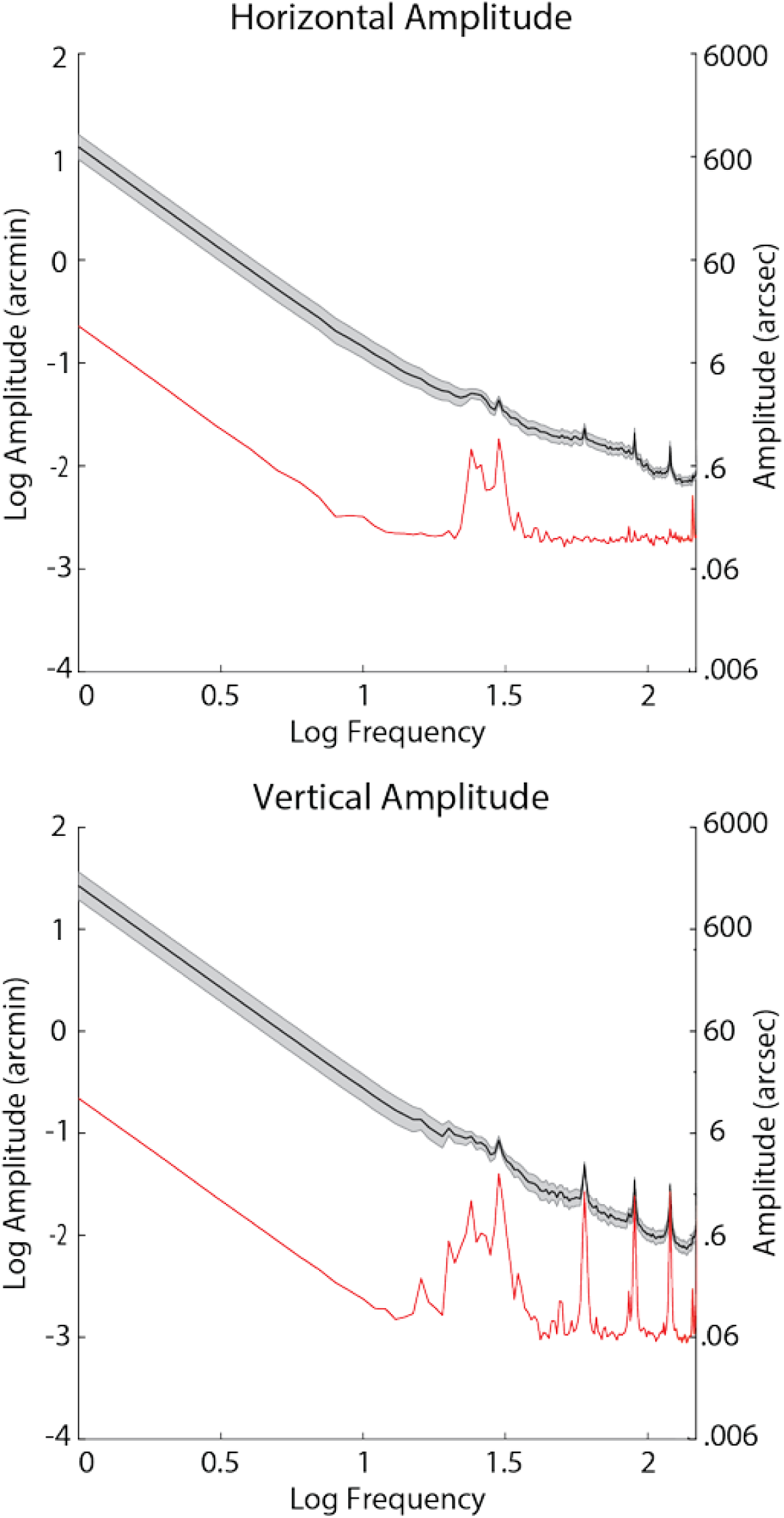
Amplitude spectra from fixational drift measured using the tracking scanning laser ophthalmoscope from 6 subjects (black) and noise floor measured from a non-moving model eye (red). Eye motion with or without cycloplegia show a similar lack of tremor on the retinal image.

### Why are the current measurements lower than all previous reports?

At first glance, the negligible increase in the amplitude spectrum indicative of tremor on the retinal image is a surprising finding considering previous reports. The statistics of tremor vary greatly across different studies, with reported amplitudes as small as 4.8 arcsec and as large as 1 arcmin (Ko, et al., 2016; Ratliff & Riggs, 1950). Ultimately, tremor is not a well-defined concept and there appears to be large differences across subjects, eye tracking techniques, and quantitative analyses. Nevertheless, tremor has been consistently observed in eye motion traces obtained from a number of high-resolution eye tracking systems. It is important to note however, that all reports of tremor to date have relied on tracking eye motion from the anterior segment of the eye, and retinal image motion has only ever been inferred. The current study is based on unambiguous, high-resolution measurements of the retinal image motion directly.

Here, we describe a combination of optical and biophysical factors that may serve to reduce the amplitude of tremor of the retinal image relative to entire eyeball. Some evidence indicating the presence of such reduction can be found in a report where AOSLO and DPI traces were recorded simultaneously (Stevenson & Roorda, 2005). In that report, eye motion traces from the two modalities were very similar except after microsaccades. However, no effort to compare estimates of tremor between the two tracking modalities was attempted in that paper. They speculated that overshoots caused by lens wobble that were detected in the DPI trace resulted in much smaller overshoots in movement of the retinal image as recorded in the AOSLO traces. Their experimental finding confirmed predictions made by Deubel and Bridgeman (1995).

There are two stages to modeling the differences between eye motion measured from the anterior segment and eye motion measured from retinal images. First is optical modeling and second is an analysis of the temporal dynamics of the lens.

### Optical Modeling

We used optical design software (Zemax, LLC, Kirkland Washington, USA) to model the effects of lens tilt and decentration in the schematic eye model available from the Zemex website (http://customers.zemax.com/os/resources/learn/knowledgebase/zemax-models-of-the-human-eye). Based on the optical model, we found that lens displacements and tilts both give rise to retinal image motion (Figure 7). Lens translations have the most profound effect: laterally displacing the lens by 1 mm gives rise to a 0.95 degree displacement of the retinal image (as measured from the second nodal point of the eye) in the same direction. Tilting the lens also displaces the retinal image: the retinal image moves in the same direction as the movement of the optical axis of the lens, but the magnitude is negligible compared to displacement. There is a −0.042 deg movement of the retinal image (measured from the second nodal point) per degree eye of lens rotation.

**Figure 7.**
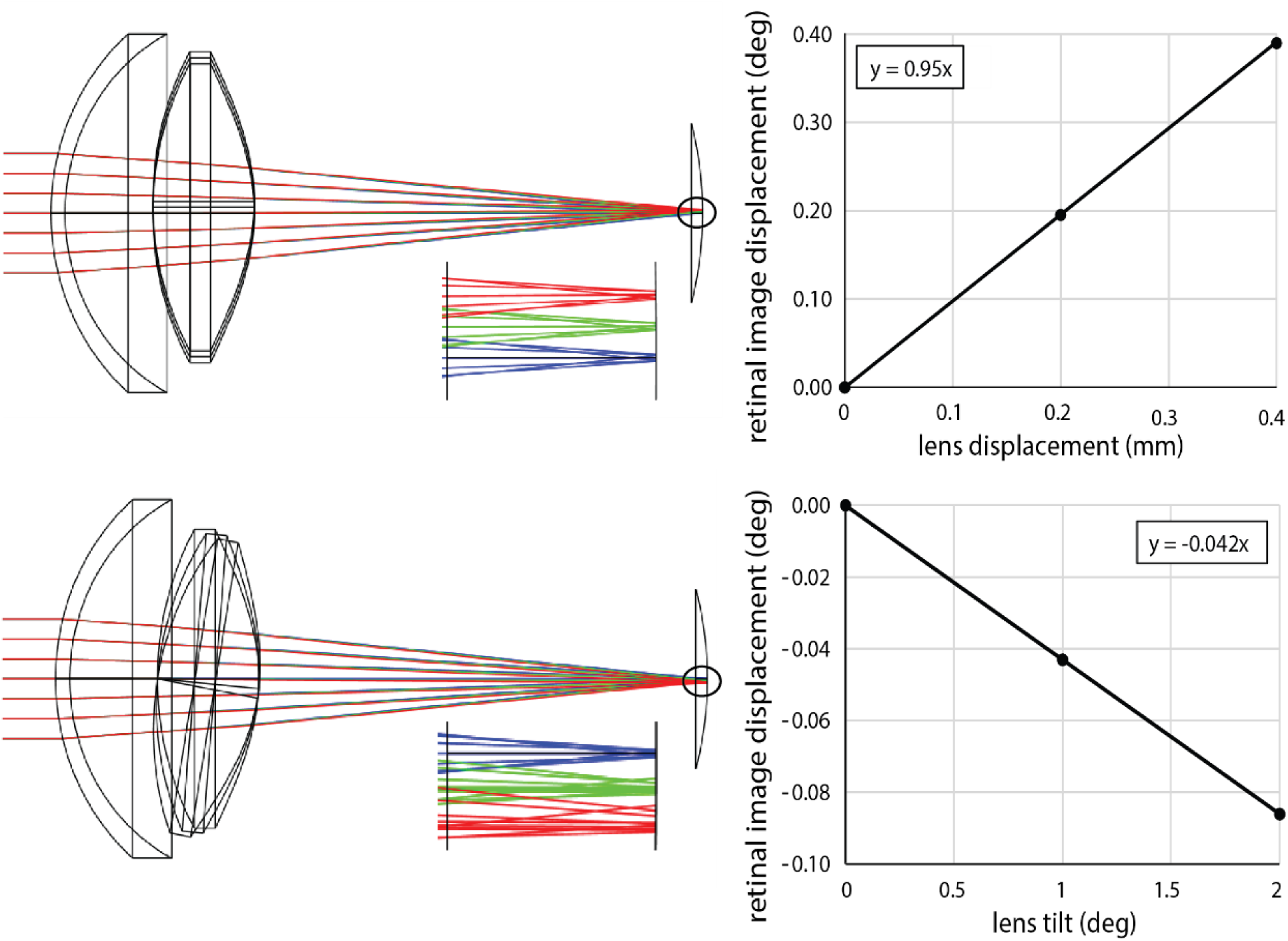
Zemax simulations. Top. Lens displacements of 0, 0.2 and 0.4 mm are illustrated in the drawing. With lens displacement, the image is displaced in the same direction. 1 mm of lens displacement gives rise to just under 1 deg of displacement of the retinal image (assuming that 1 deg visual angle corresponds to 300 microns on the retina). Bottom. Lens tilts of 1 and 2 deg were tested (amplified tilts of 5 and 10 deg are shown in the schematic to help to visualize the effect). In this case, the image displaces in the same angular direction as the tilt, but the effect is very small. 1 deg of tilt gives rise to −0.043 degs of retinal image displacement.

### Temporal Dynamics of the Lens

We considered how the lens might move within the tremoring eye, owing to the fact that it is elastically supported by the zonules and ciliary body. The manner in which the crystalline lens moves in the eyeball following a saccade is a classic example of <u>damped harmonic motion</u>. He et al. (2010) confirmed this to be the case when they measured the motion of the lens following a saccade. A frequency analysis of one of the subjects in the study revealed the resonant frequency of lens oscillation to be about 20 Hz. The oscillation of the lens over time, *x(t)*, dampens quickly, after about 1-2 cycles, following the equation:

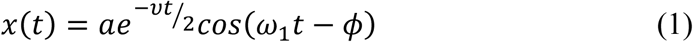

where *ω*_*1*_ is the oscillating frequency, given by

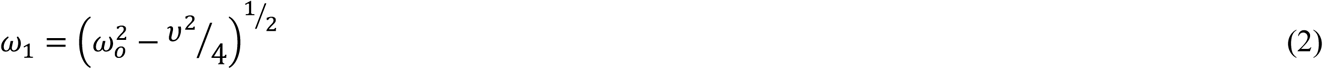

*ω*_*o*_ is the resonant frequency, *a* is the amplitude, *φ* is the phase offset (not very important), and *ν* is a constant with dimensions of angular frequency indicating the strength of damping.

Based on visual observation of the lens wobble artifact in He et al. (2010), we estimated the constant *ν* to be about ½ of the resonant frequency. A damped oscillation with a constant *ν* = *ω*_*0*_ */2* is plotted on Figure 8A and shows that the oscillation relaxes after one or two cycles.

**Figure 8.**
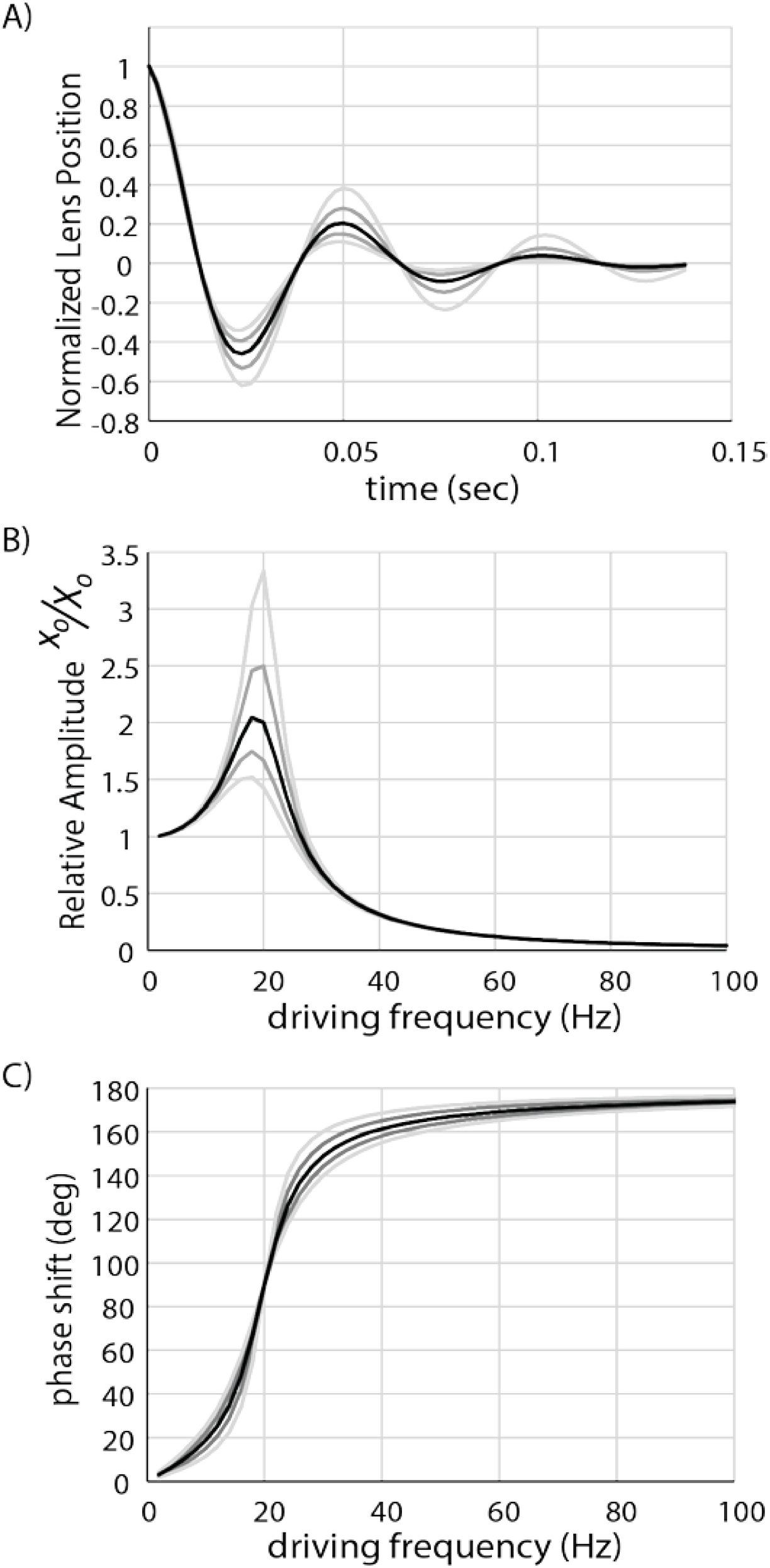
A) Model of the position of a lens behaving as damped harmonic oscillator. In this case, the damping coefficient is 20π (10 Hz), or half of the resonant frequency of 20 Hz, which leads to about two cycles of oscillation before relaxing to zero. Gray shaded lines in all three plots show calculations with +/-20% and +/-40% changes in the damping coefficient. B) Model of the amplitude of a lens behaving as a driven damped harmonic oscillator with resonant frequency of 20 Hz and range of damping coefficients. Driving frequencies near the resonant frequency gives rise to amplified motion of the lens. Driving frequencies within the range of tremor give rise to lens oscillations that are, on average, about 0.1 of the driving amplitude. Amplitudes of lens motion at the resonant frequency are highly dependent on the damping coefficient, but outside of that the trends remains relatively similar. C) Model of the phase shift of the lens motion. When driving frequencies are slow, the lens moves along with the eyeball as expected. For driving frequencies that are in the range of tremor (50-100 Hz) the lens moves in counter-phase with the eyeball. Changes in the damping coefficient have little effect on this trend.

The manner in which tremor affects the motion of the lens is a classic example of <u>driven damped harmonic oscillation</u>. The eyeball rotates about its center of rotation, and how the lens reacts to this motion depends on the damping constant *ν* and the resonant frequency *ω*_*o*_ in the following way.

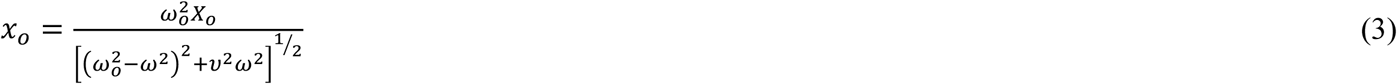

and

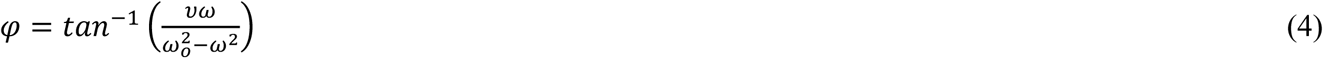

where *x*_*o*_ is the amplitude of the oscillating mass (the lens), *X*_*o*_ is the amplitude of the driver (eye tremor), *ω*_*o*_ is the resonant frequency, *ω* is the frequency of the oscillating driving force (eye tremor) and *φ* is the phase shift between the driving oscillation and the oscillating mass.

Figure 8B plots the amplitude and phase of the lens oscillation that has a resonant frequency of *ω*_*o*_ = *40π (20 Hz)* and a damping coefficient of *ν* = *0.5ω*_*o*_ over a range of driving frequencies. Note that for low frequencies, we see the expected behavior where the lens moves in phase with the eye with the same amplitude. When the driving frequency reaches the resonant frequency of the lens, the amplitude oscillation increases to about 2 times. At the same time the phase of the lens movement transitions to counterphase motion. When the driving frequency increases further, the motion is in counterphase and the amplitude approaches zero. In the limit, the lens remains perfectly fixed in place relative to the tremoring eye. The modeled behavior remains qualitatively similar with variations in the damping constant *ν*, which were deduced from plots in He et al. (2010). Changes in the damping coefficient by up to 40% do not change the general trends in the plots. Similarly, changes in the resonant frequency will shift the curves but not in a manner that would alter the main conclusion, which is that during tremor, the lens moves with lower amplitude than the eyeball and it oscillates in counterphase to the eyeball.

A counterphase motion of the lens relative to the eyeball would, in effect, amplify the motion that is measured by the DPI eye tracker. The situation is illustrated schematically on Figure 9. In the DPI, the eye rotation is assumed to be proportional to the separation between the reflection from the cornea, P1, and the reflection from the back surface of the lens, P4, which are situated approximately at the center of curvatures of the surfaces that generate them (Cornsweet & Crane, 1973). If the lens and eyeball rotate together, as they would with slow rotations, then the separation between the two reflexes (assuming small angles) is *∼7φ* where *φ* is the rotation angle [in radians] (Cornsweet & Crane, 1973). If the lens moves in counterphase to the eyeball rotation with 1/10 of the amplitude (as per estimations on Figure 8C), then the separation is ∼ *−7.3 φ* The overall magnitude of the separation between the two reflexes is 4.2 % larger than the actual motion, and the sign is opposite.

**Figure 9.**
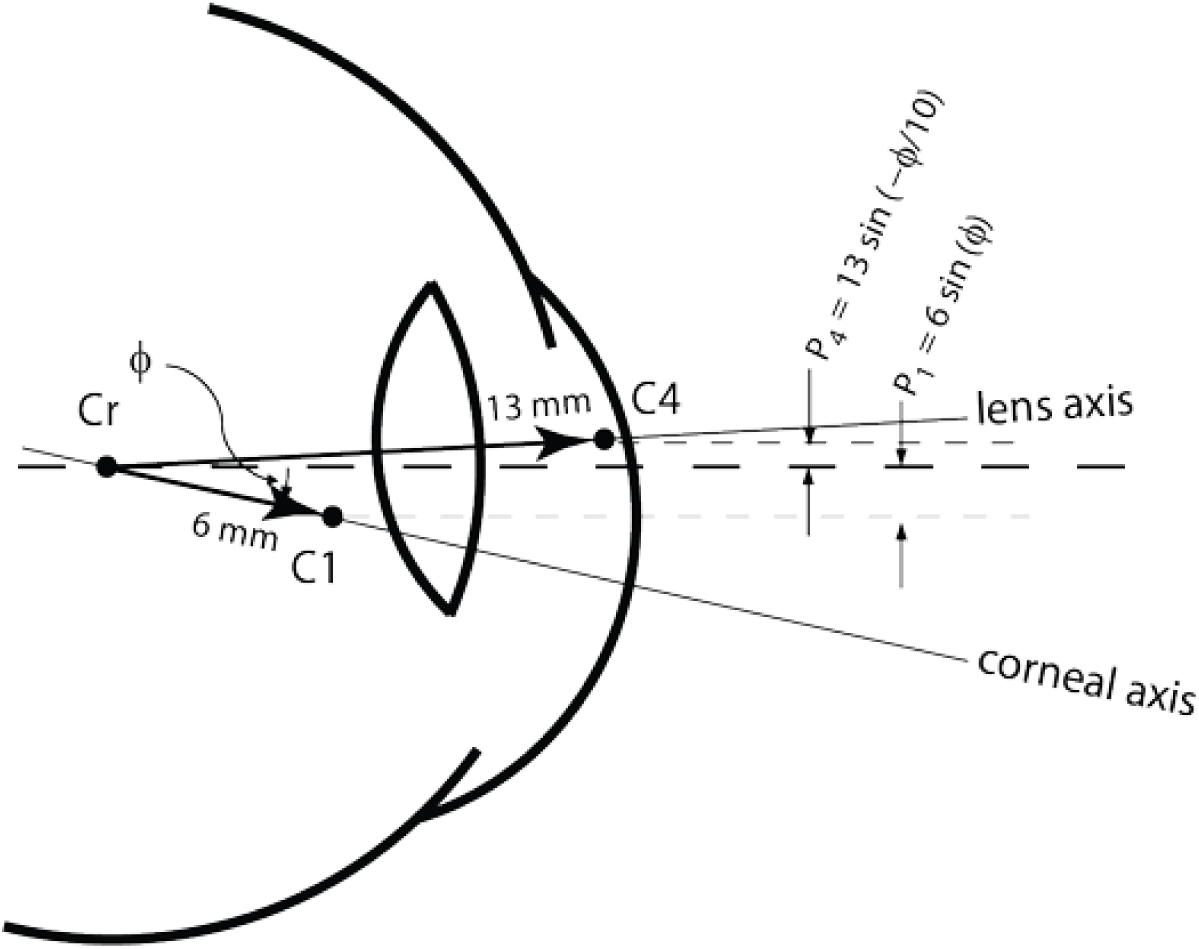
Schematic drawing showing how Purkinje reflections get displaced with rotations of the eye and lens. In the schematic, the eyeball has rotated downward by an angle φ and the lens has undergone a counterphase movement upward with 1/10 of the amplitude. For collimated incident light, the Purkinje reflections P1 and P4 are approximately positioned at the centers of curvature of their respective reflecting surfaces. The two reflections are displaced according to the equations on the figure. The total separation between C4 and C1 is 13sin(-φ/10) - 6sin(φ) which, for small angles, is −7.3 φ

The same lens translations would minimize the amplitude of the motion of the retinal image. Consider an eyeball undergoing a tremor rotational motion with an amplitude of 1 minute. The amplitude of pupil displacement associated with the rotational eye movement is about 0.0029 mm (since the pupil is displaced about 10 mm from the center of rotation of the eye, giving rise to displacement of *S*_*P*_ = *10 sin (φ)).*

The lens will move this amount plus an additional 10% due to the counterphase oscillation for a total displacement of 0.0032 mm. These lens displacements cause the amplitude of the retinal image motion to be smaller, by 0.003 deg (0.18 arcmin) or 82% of the eyeball rotation. In total, AOSLO measures of tremor amplitude should be 82% - 4.2% ≈ 78% of DPI measures.

However, tremor is not only detected in DPI traces, but has been seen in eye motion traces from search coil measurements as well as from traces of reflections from the cornea, neither of which involve a measurement of the lens. In these cases, assuming there is no artifact from the search coil or corneal reflection measurement, the only effect that will diminish the movement of the image on the retina from the overall eyeball motion is the translation of the lens which, according to the Zemax model, will give rise to a retinal motion amplitude that is diminished by ∼ 18%.

Considering the arguments made above, the reported tremor amplitudes of 4.8 and 6 arcsec from Ko et al. (2016) and Eizenman (1985), respectively (Eizenman, et al., 1985; Ko, et al., 2016), would reduce to 3.75 and 4.92 arcsec tremor amplitude on the retina. Our results are still lower than these previous reports, even after correction, but they are in the same order of magnitude. The remaining differences could be due to the actual fixation task: our subjects were actively engaged in a fixation/acuity task, but Ko et al. had their subjects simply fixate the center of a blank screen and Eizenman et al. had their subjects fixate on a small source at 50 cm. Additionally, other biophysical factors, such as damping of lateral motion due to the elasticity of the retinal surface, could also be present. More experiments would be required to assess the effects of these possible causes for the differences.

## Conclusion

In this study we validated the use of the AOSLO as a high-resolution eye tracking technique that is uniquely able to resolve small movements on the retinal image. We first validated the capabilities of the AOSLO system by recording movies of an oscillating model eye and reliably recovering both the frequency and amplitude put into the galvanometers. We also were able to recover sinusoidal motion that was artificially inserted into a real AOSLO movie. The AOSLO system was able to reliably recover the fixational eye motion of 6 human subjects as they participated in a tumbling E task. The statistics of both microsaccades and fixational drift were generally found to be within normal parameters of previous reports using high-resolution eye tracking. However, the amplitude of fixational tremor was smaller than all previous reports. Although the true cause of this reduction of tremor on the retinal image is beyond the scope of the paper, we suggest an optical model relying on recent reports that the lens can move independently of the eye that may partially account for the loss of power within this frequency band.

## Acknowledgements

This work was supported by NEI NIH grants R01EY023591, P30EY001730 & T32EY007043. As well as the Minnie Flaura Turner Memorial Fund for Impaired Vision Research and the Michael G. Harris Ezell Fellowship.

